# A mechanistic understanding of the modes of Ca ion binding to the SARS-CoV-1 fusion peptide and their role in the dynamics of host membrane penetration

**DOI:** 10.1101/2022.03.03.482731

**Authors:** Juliana Debrito Carten, George Khelashvili, Miya K. Bidon, Marco R. Straus, Tiffany Tang, Javier A. Jaimes, Harel Weinstein, Gary R. Whittaker, Susan Daniel

**Affiliations:** Robert Frederick Smith School of Chemical and Biomolecular Engineering, Cornell University, Ithaca, NY, USA; Departments of Microbiology & Immunology and; Public & Ecosystem Health, Cornell University, Ithaca, NY, USA; Department of Physiology & Biophysics, Weill Cornell Medicine, New York, NY, USA; Institute for Computational Biomedicine, Weill Cornell Medicine, New York, NY 10065, USA

## Abstract

The SARS-CoV-1 spike glycoprotein contains a fusion peptide (FP) segment that mediates fusion of the viral and host cell membranes. Calcium ions are thought to position the FP optimally for membrane insertion by interacting with negatively charged residues in this segment (E801, D802, D812, E821, D825, and D830); however, which residues bind to calcium and in what combinations supportive of membrane insertion are unknown. Using biological assays and molecular dynamics studies, we have determined the functional configurations of FP-Ca^+2^ binding which promote membrane insertion. We first mutated the negatively charged residues in the SARS CoV-1 FP to assay their role in cell entry and syncytia formation, finding that charge loss in the D802A or D830A mutants reduced syncytia formation and pseudoparticle transduction. Interestingly, the D812A mutation led to increased pseudoparticle transduction, indicating the Ca^2+^ effect depends on binding at specific FP sites. To interpret mechanistically these results and learn how specific modes of FP-Ca^2+^ binding modulate membrane insertion, we performed molecular dynamics simulations. Preferred residue pairs for Ca^2+^ binding were identified (E801/D802; E801/D830; D812/E821) which promote FP membrane insertion. In contrast, binding to residues E821/D825 inhibited FP membrane insertion, which is also supported by our biological assays. Our findings show that Ca^2+^ binding to SARS-CoV-1 FP residue pairs E801/D802 and D812/E821 facilitates membrane insertion, whereas binding to the E801/D802 and D821/D825 pairs is detrimental. These conclusions provide an improved and nuanced mechanistic understanding of calcium binding modes to FP residues and their dynamic effects on host cell entry.

---

Coronaviruses (CoVs) are a diverse family of enveloped single-stranded RNA viruses responsible for causing respiratory and enteric diseases across a wide range of species (1, 2). Three human coronaviruses have posed significant public health threats during the early 21^st^ century: the severe acute respiratory syndrome coronavirus (SARS-CoV-1) in 2002, (3), Middle East respiratory syndrome coronavirus (MERS-CoV) in 2012 (4), and currently SARS-CoV-2 (5, 6). SARS-CoV-2 vaccines have proven to be highly effective at preventing morbidity and death during the pandemic; however, current experience demonstrates that there is an ever-present need for antivirals as a parallel line of defense. Development of such therapeutics requires a better understanding of the fundamental biology of coronaviruses and the intrinsic and extrinsic determinants of viral entry.

Infectivity by coronaviruses is mediated by the highly conserved and complex spike (S) glycoproteins, which form homotrimers in the virion membrane and interact with host cell receptors, such as angiotensin-converting enzyme 2 (ACE2) or DPP4 (7) during the entry process. Upon spike-mediated interaction with the host cell surface receptor, CoVs are proposed to utilize two different entry pathways: an “early” route involving proteolytic priming at the cell surface and a “late” route that follows the canonical endosomal/lysosomal pathways (8). The entry pathway utilized is largely dependent on the local protease environment (9, 10). Previous work has shown that SARS-CoV-1 can enter host cells via the early pathway when the transmembrane serine protease 2 (TMPRSS2) and the virus receptor ACE2 are both present in the plasma membrane, or when activated by exogenous trypsin after binding to the cell surface (11). SARS-CoV-1 can also enter cells via the endosomal route where low pH-dependent proteases, such as cathepsin L, cleave and prime the FP for membrane insertion and fusion. This latter pathway is likely the predominant one used by SARS-CoV-1 in most laboratory cell lines used thus far (e.g. Vero cells)(9,10,12).

The entry flexibility seen across coronaviruses arises from the differential requirements for protease priming and activation of the S protein. The SARS-CoV-1 S protein contains proteolytic cleavage sites that enable its spatial and temporal regulation throughout the viral life cycle (9). Two well-characterized protease sites, S1/S2 (R667) and S2’ (R797), are specifically involved in SARS-CoV-1 FP activation. The S1/S2 site resides at the boundary between the S1 (receptor binding) and S2 (fusion) domains of the spike and is cleaved by trypsin-like proteases or cathepsin (9, 13, 14). The S2’ prime site is located adjacent to the N-terminus of the FP and cleavage at this residue allows the S2 domain to undergo a large conformational change that favorably positions the FP for insertion into the host cell membrane (12).

In addition to protease priming, it is known that several viral fusion glycoproteins need calcium ions to initiate membrane fusion during viral entry. The Rubella virus spike glycoprotein (E1) was the first calcium-dependent viral fusogen discovered, with calcium found to play a role specifically on E1 membrane insertion and fusion (15). Subsequent studies done by our group with Zaire Ebolavirus (16), SARS-CoV-1 (17), and MERS-CoV (18) have shown that calcium interacts directly with viral fusogens and is a conserved feature across several different viral entry pathways. In work done with the SARS-CoV-1 and MERS-CoV FPs, calcium was also found to induce membrane ordering in a calcium-dependent manner following FP membrane insertion, with the stoichiometry between the FP segments and calcium ions predicted to be 1:2 for SARS-CoV-1 and 1:1 for MERS-CoV (18). Additionally, the SARS-CoV-1 FP adopts a higher degree of α-helicity in the presence of calcium ions and synthetic membranes (17, 19). Taken together, these results suggest that direct interaction between calcium ions and the SARS-CoV-1 FP residues stabilizes the spike FP to help anchor it in the host cell membrane to promote viral-host membrane fusion.

The CoV FP sequence contains several conserved negatively charged amino acids that flank the highly conserved hydrophobic residue motif LLF, which is known to interact with the host cell membrane upon FP insertion (17, 20). We have posited that, like the SARS-CoV-2 FP (21), the SARS-CoV-1 FP binds to calcium at these negatively-charged residues to stabilize its local structure and position it for optimal membrane penetration (17, 19, 21). However, we lack a detailed mechanistic understanding of how binding modes of Ca^2+^ to the SARS-CoV-1 FP charged residues facilitates host membrane insertion. The negatively charged residues of the SARS-CoV-1 FP important for calcium binding were identified from a combination of mutagenesis studies (22) and biophysical characterizations (23), which found that when mutated, these residues’ interactions with calcium were affected to varying degrees.

Here we first systematically examine the effects of mutating charged residues in the SARS-CoV-1 FP on functional metrics consisting of syncytia formation and pseudoparticle transduction, two assays which assess membrane fusion and entry, in cultured cells. We chose SARS-CoV-1 as a prototype beta CoV model FP as it has the best prior analysis of structure-function relationships. We then used atomistic molecular dynamics (MD) simulations of the corresponding SARS-CoV-1 FP molecular systems to determine where Ca^2+^ binds preferentially and how these modes of binding affect FP membrane insertion. The analysis of the preferred loci of Ca^2+^ binding for the wild type and mutated FP segments identified two residue pairs where Ca^2+^ binding was observed to promote FP membrane insertion (E801/D802; D812/E821), and one pair where Ca^2+^ binding inhibits insertion (E821/D825). Experimental probing using biological assays largely confirmed these predictions. Our interpretations of these results enabled us to propose two modes of SARS-CoV-1 FP membrane penetration modulated by FP-calcium interactions: (Mode 1) Binding of Ca to FP residue pairs E801/D802 and D812/E821 supports insertion of the LLF-containing segment by shielding the neighboring negative charges from repulsion by the phospholipid headgroups, and (Mode 2) Binding of Ca to FP residue pair D821/D825 stabilizes a conformation that leaves unshielded anionic residues in the LLF-containing insertion region, which is unfavorable for membrane penetration. These mechanistic inferences from our combined biological and computational investigation provide an atomistically nuanced understanding of how specific FP residues interact with calcium ions to drive host membrane insertion during SARS-CoV-1 infection.

## Results

### Generation and characterization of SARS-CoV-1 Spike fusion peptide mutants

Various structural conformations of the SARS-CoV-1 spike monomer have been determined using Cryo-EM (PDB: 5XLR, 5WRG, 5X58) (24). In these pre-fusion structures, the fusion peptide (FP) segment containing the core hydrophobic residues LLF appears to adopt an α-helical conformation, followed by a proximal disordered loop known to participate in cysteine-mediated intramolecular disulfide bonding (**Fig 1A**). The 6 negatively charged residues (E801, D802, D812, E821, D825, D830) in this SARS-CoV-1 FP segment are highly conserved across SARS-CoV-1, MERS-CoV, and SARS-CoV-2 (**Fig 1B**).

**Figure 1.**
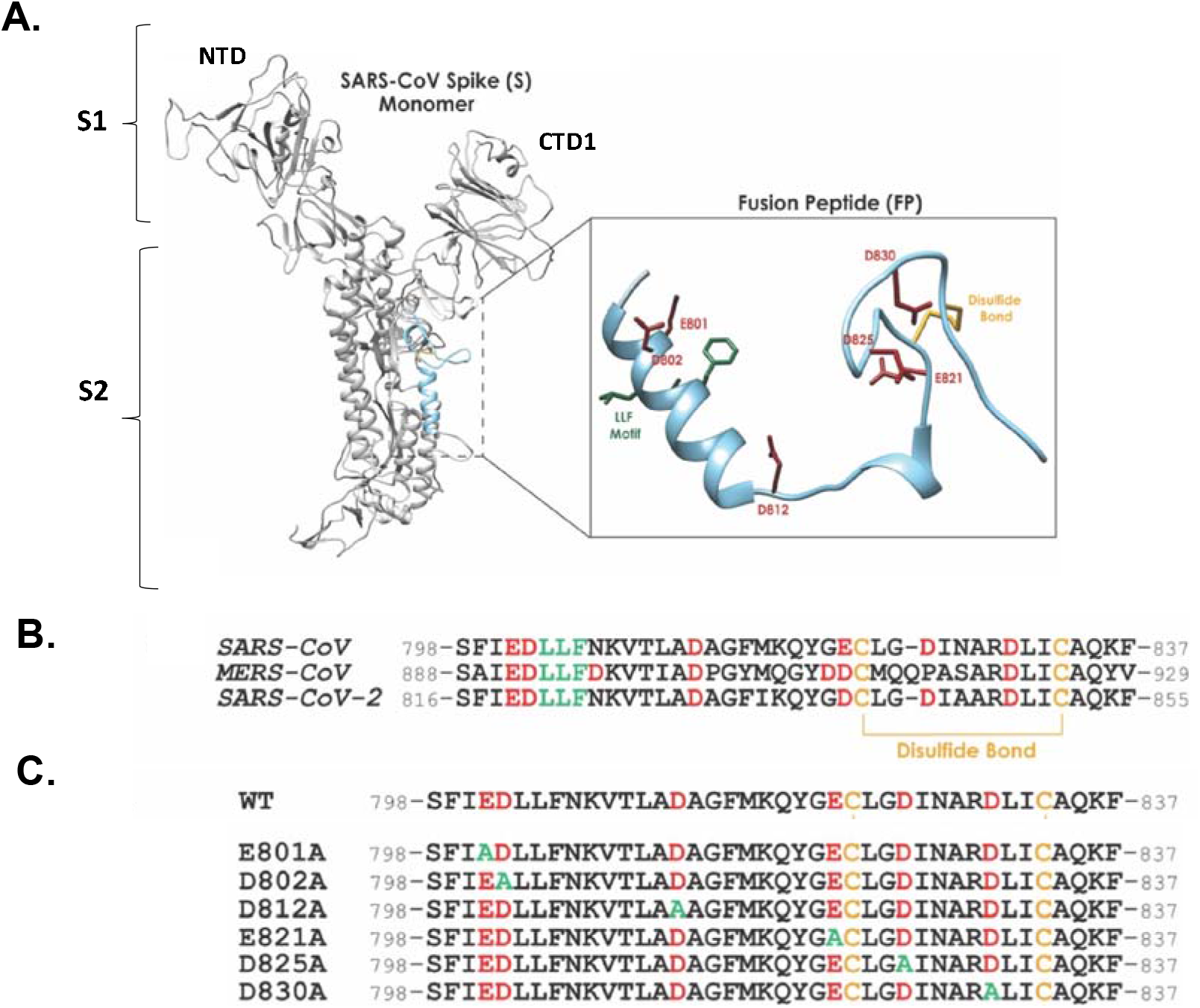
Structure of WT SARS-CoV-1 spike monomer, the fusion peptide region, and the amino acid sequences of the S constructs generated in this study. (A) The monomeric SARS-CoV-1 spike glycoprotein in the prefusion conformation (Gui et al. 2017; PDB: 5XLR). The β strand-rich S1 subunit of the spike protein contains the C-terminal domain 1 (CTD1), which mediates ACE2 receptor binding, and the N-terminal domain (NTD). The S2 subdomain contains mostly α-helices and is comprised of two heptad repeat regions (HR1 and HR2) and the fusion peptide (FP). The fusion peptide is highlighted in light blue and expanded to show its predicted structure and residues of interest in this study. The hydrophobic LLF residues appear in green, negatively charged residues in red, and the disulfide bond in yellow. (B) Alignment of the amino acid sequences of SARS-CoV-1, MERS-CoV, and SARS-CoV-2 FPs showing the hydrophobic LLF motif (green), the negatively charged residues (red), and the intramolecular disulfide bond. (C) The 40-amino acid long sequence of the SARS-CoV-1 WT FP (GenBank accession no. AAT74874.1). Charged residues appear in red, alanine substitutions appear in green, and cysteine residues appear in yellow.

To determine the functional roles of these negatively charged residues and their involvement in the binding of Ca^2+^ during viral entry, we generated single and double alanine substitutions of these residues. We first made single amino acid (aa) substitutions (E801A, D802A, D812A, E821A, D825A, D830A) in the wild-type SARS-CoV-1 FP. The wildtype and mutant forms of the SARS-CoV-1 S gene were then cloned into pcDNA3.1 expression vectors to enable their transient expression and characterization. To verify the synthesis of folded and active protein, we assessed the ability of the spike (S) protein to be appropriately cleaved for fusion activation. To achieve this, 24 hours after transfection, HEK293T cells were incubated with TPCK-trypsin (1ug/mL) for 10 minutes. Cell surface S proteins were biotinylated and retrieved following cell lysis with streptavidin beads, resolved by gel electrophoresis, and immunoblotted for using a SARS-CoV-1 S polyclonal antibody. The resulting spike immunoblots displayed the full-length, uncleaved S species (S_0_) migrating at ∼185kDa in untreated samples (**SI Fig S1 A, odd lanes**). SARS-CoV-1 S mutants containing the single mutations D802A, E821A, D825A, and D830A had comparable cell surface levels of S protein to the wild-type protein, indicating that these mutants are not impaired in their synthesis or trafficking to the plasma membrane. Additionally, proteolytic cleavage of all mutants occurred following treatment with Trypsin, indicating that these mutants had accessible cleavage sites and were able to be primed for downstream fusion evens (**SI Fig S1 A, even lanes**). The two lower molecular weight bands that appear at ∼100kDa and ∼80kDa in the TPCK-trypsin treated lanes are the S_1_ and S_2_ subunits, which are generated following cleavage presumably at the S1/S2 site (**SI Fig S1,** black arrows). These subunits have been detected previously following cleavage of a recombinant S protein monomer by Trypsin at the S1/S2 cleavage site (25). We note that we were unable to detect surface levels of the E801A mutant on the spike immunoblots (data not shown). To further probe the nature of this residue’s importance for S protein stability, we substituted it with the larger, nonpolar methionine (E801M), a polar and uncharged glutamine (E801Q), a positively charged lysine (E801K) and the charge mimetic aspartic acid (E801D). None of these mutations restored the cell surface levels of the S protein to that of wild-type (data not shown), indicating the specific requirement of glutamic acid in this region of the FP.

### Analysis of the SARS-CoV-1 FP single mutant pseudoparticles’ ability to transduce cells

We next tested the ability of the SARS-CoV-1 single mutants to facilitate host cell entry using SARS-CoV-1 pseudoparticles as a surrogate for infectious virus. This was necessary because SARS-CoV-1 is designated a Risk Group 3 Select Agent which requires experiments to be carried out in a specialized biosafety level 3 (BSL-3) facility. The pseudoparticles offer a safe alternative to live SARS-CoV-1 virus work and enable experiments to be done in BSL-2 level conditions (26). To generate SARS-CoV-1 pseudoparticles in HEK293T cells, we used a three-plasmid co-transfection system with plasmids encoding: (1) the full length CoV-1 surface S protein, (2) the murine leukemia virus (MLV) core proteins gag and pol, or (3) a firefly luciferase reporter gene containing the MLV-MJ RNA packaging signal and flanked by long terminal repeat (LTR) sequences (26). Upon successful pseudoparticle fusion with the host cell membrane, the luciferase reporter RNA transcript is reverse transcribed, integrated into the cellular genome, and expressed – which enables a measurable readout of pseudoparticle entry via luciferase activity. Our previous studies have shown that this approach works well for assessing the infectivity of MERS-CoV (18), SARS-CoV-1 (17, 26), and SARS-CoV-2 (27) pseudoparticles.

Incorporation of the WT and mutant S proteins into the SARS-CoV-1 pseudoparticles was assessed via Spike immunoblotting of the harvested pseudoparticles. Pseudoparticles with WT or the mutant S proteins D802A, D812A, E821A, D825A, and D830A displayed S protein incorporation, with bands at ∼185 kDa (S_0_) representing full-length monomeric S protein appearing in all lanes (Fig 2A).

**Figure 2.**
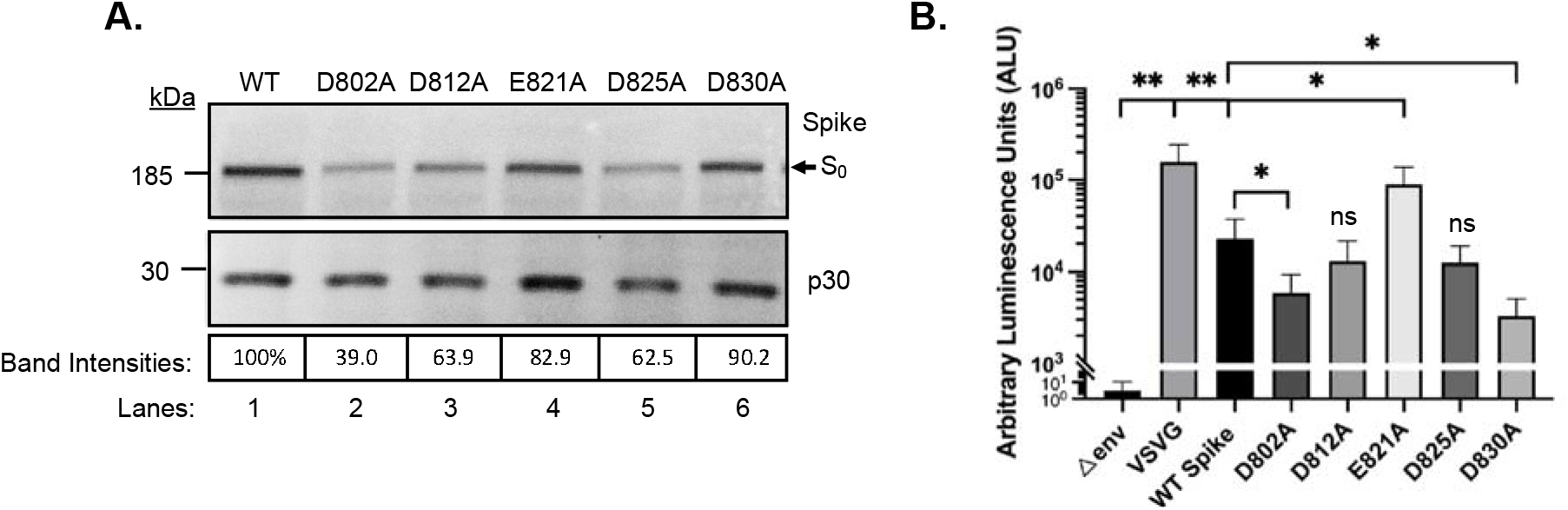
Immunoblot analysis and transduction assays of wild-type and single mutant SARS-CoV Spike-containing pseudoparticles. (A) Representative Spike and p30 immunoblots of pseudoparticles containing wild-type (WT, lane 1) or single charge-to-alanine spike mutants (lanes 2 to 6). S_0_ indicates the full-length, uncleaved spike protein. MLV p30 was used as a loading control. Spike and p30 band intensities from 2 sets of blots done with a single batch of pseudoparticles were quantified using ImageJ. Band intensities were normalized to WT and appear as percentages below the blot lanes. (B) WT and mutant SARS-CoV-1 pseudoparticle transduction assays in VeroE6 cells. VSVG-typed and delta envelope (Δenv) pseudoparticles were used as positive and negative controls respectively. * and ** denote a significance of p ≤ 0.05 or p ≤ 0.01, respectively. Graph represents 5 technical replicates of the same batch of pseudoparticles. The transduction assay was repeated with a second batch of pseudoparticles with a minimum of 3 technical replicates.

Spike band intensities were quantified relative to the loading control (p30) and then normalized to the WT spike band, which was set to 100%. All of the single mutant-containing pseudoparticles had reduced levels of full-length spike to varying degrees (**Fig. 2A**). The D802A mutant had the least amount of full-length Spike (39.0%), followed by the D812A (63.9%) and D825A (62.5%) mutants. For the E821A and D830A mutants we measured, only minor decreases in Spike levels were observed at 82.9% and 90.2%, respectively. This variability in fusogen levels across the pseudoparticles generated was considered when interpreting the results of the transduction assay. After confirming the incorporation of the spike proteins into the SARS-CoV-1 pseudoparticles, we tested their ability to enter permissive cells. Exposure of VeroE6 cells to WT SARS-CoV-1 pseudoparticles results in a robust luminescent signal 72 hours post transduction, indicating successful pseudoparticle and cell membrane fusion (**Fig 2B**). Pseudoparticles containing the D812A and D825A Spike mutants exhibited luminescence values comparable to WT levels, indicating that these individual mutations do not affect cell entry despite containing ∼63% of the WT levels of Spike (**Fig 2A, lanes 3 and 5**). In comparison, the D802A and D830A mutant pseudoparticles showed a decrease in cell transduction, resulting in 3.8- or 6.8-fold reduction in luminescence values, respectively. It should be noted that the D802A pseudoparticles had the greatest reduction in Spike levels (39%), which may have contributed to the low luminescence values we observed in the transduction assay. Somewhat surprisingly, the E821A-containing pseudoparticles showed a 3.9-fold increase in luminescence, suggesting that this FP mutation enhances cell entry.

### Analysis of the SARS-CoV-1 FP mutants’ ability to promote syncytia formation

Functional tests of the ability of S mutants to induce cell-cell fusion were performed using a standard technique in the field known as a syncytia assay. Syncytia are multinucleated cells that form following the fusion of neighboring cells during SARS-CoV-1 infection and are one of several pathological hallmarks of cellular dysfunction induced by viral infection(28, 29). To perform this assay, WT or mutated S constructs were transiently transfected into VeroE6 cells, a kidney epithelial cell line that expresses the ACE2 receptor (30). 24 hours following transfection, cells were treated with TPCK-Trypsin (2ug/mL) to cleave the FP at the S1/S2 site and induce cell-cell fusion. Syncytia were visualized by immunofluorescence using a SARS-CoV-1 S antibody and cells were co-stained with DAPI to identify multinucleated cells. Cells transfected with WT SARS-CoV-Spike and left untreated served as a negative control for syncytia formation. Syncytia were quantified by counting every group of fused cells with a minimum of 4 nuclei and expressed as a percentage of the total number of generic fluorescent particles counted in a field of view (yes/no binary binning method). VeroE6 cells expressing the WT S readily formed syncytia with an average of ∼45.1% of fluorescent regions of interest (ROIs) being classified as syncytia (**Fig 3**). Similarly, VeroE6 cells expressing the E821A (40.3 % syncytia) or D825A (43.7% syncytia) Spike mutants exhibited comparable syncytia formation to cells expressing WT S, indicating that loss of a negative charge in these individual residues does not affect FP function (**Fig 3**). Conversely, cells expressing the D802A, D812A, and D830A Spike show defective syncytia formation, with the D802A (13.5 % syncytia) and D830A (9.8 % syncytia) mutants showing a major reduction in syncytia formation, similar to the untrypsinized controls (9.3% syncytia). The D812A mutation moderately decreased syncytia formation to 34%, indicating that this FP residue is less important in mediating cell-cell fusion events.

**Figure 3.**
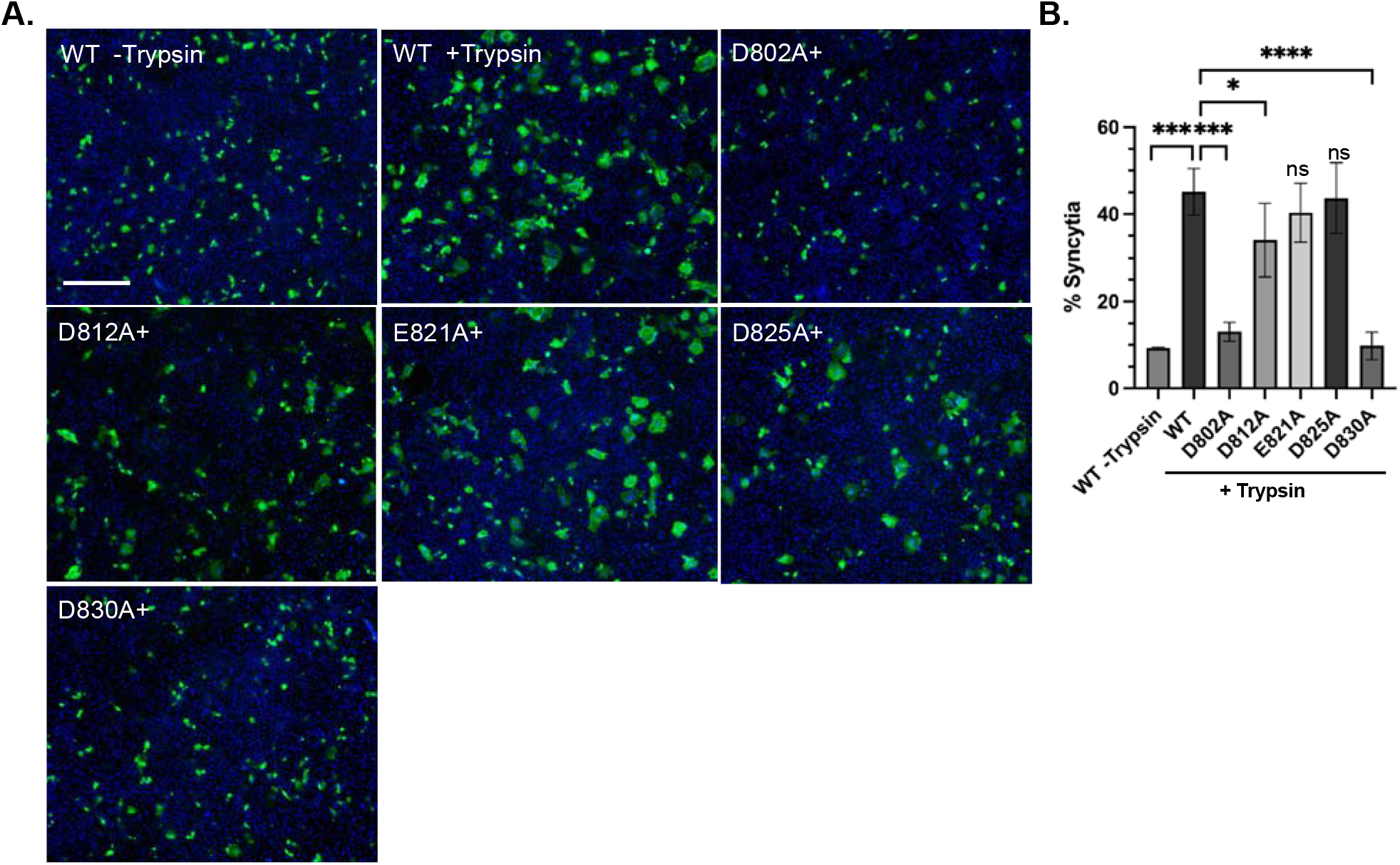
Syncytia formation in SARS-CoV-1 wild-type and single charge-to-alanine mutant S-expressing VeroE6 cells. (A) Representative immunofluorescence images of VeroE6 cells expressing WT or single charge-to-alanine mutant S constructs. 24 h following transfection, cells were left untreated (-Trypsin) or treated with 2ug/mL TPCK-Trypsin (+ Trypsin or +) for 5 minutes to cleave S proteins and induce syncytia formation. Cells were then fixed the following day and syncytia were visualized using a SARS-CoV-1 S antibody (green). Nuclei were stained with DAPI and appear in blue. Images were taken at 20X magnification. Scale bars = 90 µm (B) Quantification of syncytia across each condition. *, ***, and **** denote a significance of p <0.05, 0.001, and 0.0001, respectively. n = 3 to 6 biological replicates.

### Molecular Dynamics simulations identify the modes of the SARS-CoV-1 FP’s calcium binding

The major effects on cell-cell fusion and cellular transduction observed for the mutant SARS-CoV-Spike FP constructs confirmed their importance for viral entry processes at the level of membrane fusion. To test the relationship between the preferred modes of FP residue-calcium binding and membrane insertion, we carried out atomistic molecular dynamics (MD) simulations of the various SARS-CoV-1 FP constructs in the presence of Ca^2+^ ions. This enabled us to systematically compare the mutagenesis results and deduce a potential mechanism of FP membrane insertion. Notably, this also allowed us to examine the mutation of residue E801, which could not be included in the experimental investigations, as described above.

To monitor the spontaneous binding of calcium to the FP, we collected 18 independent MD trajectories of 640ns in length for each construct. Following the analysis protocols described recently for SARS-CoV-2 FP simulations (21, 31), the various modes of interactions between the SARS-CoV-1 FP and calcium ions were assessed in the trajectories by monitoring: i) the distances between the calcium ions in solution and the side chains of all acidic residues in the peptide; and ii) the pairwise distances between the side chains of all acidic residues. Summarized in **SI Figs S2 and S3** are the observed events of simultaneous association of two calcium ions with various pairs of FP residues in the individual trajectories of the WT and the mutant systems (see red and blue rectangles). The combined statistics for each construct (i.e., the total number of binding events for different pairs of residues) are summarized in **Fig 4**. These results show the modes of calcium coordination in the WT protein, identifying the most frequent modes involving residue pairs E821/D825 (5 out of 18); E801/D802 and E801/D830 (4/18 each); and D812/E821 (2/18). Notably, a similar pattern of coordination preference was observed in our recent computational studies of calcium association with SARS-CoV-2 FP (21). Moreover, simultaneous binding of calcium ions to the SARS-CoV-2 FP residue pairs equivalent to E801/D802 and D812/E821 produced the peptide conformations suitable for membrane penetration. In contrast, conformations that stabilized calcium binding to residues equivalent to the E821/D825 pair did not enable sustained bilayer insertion.

**Figure 4.**
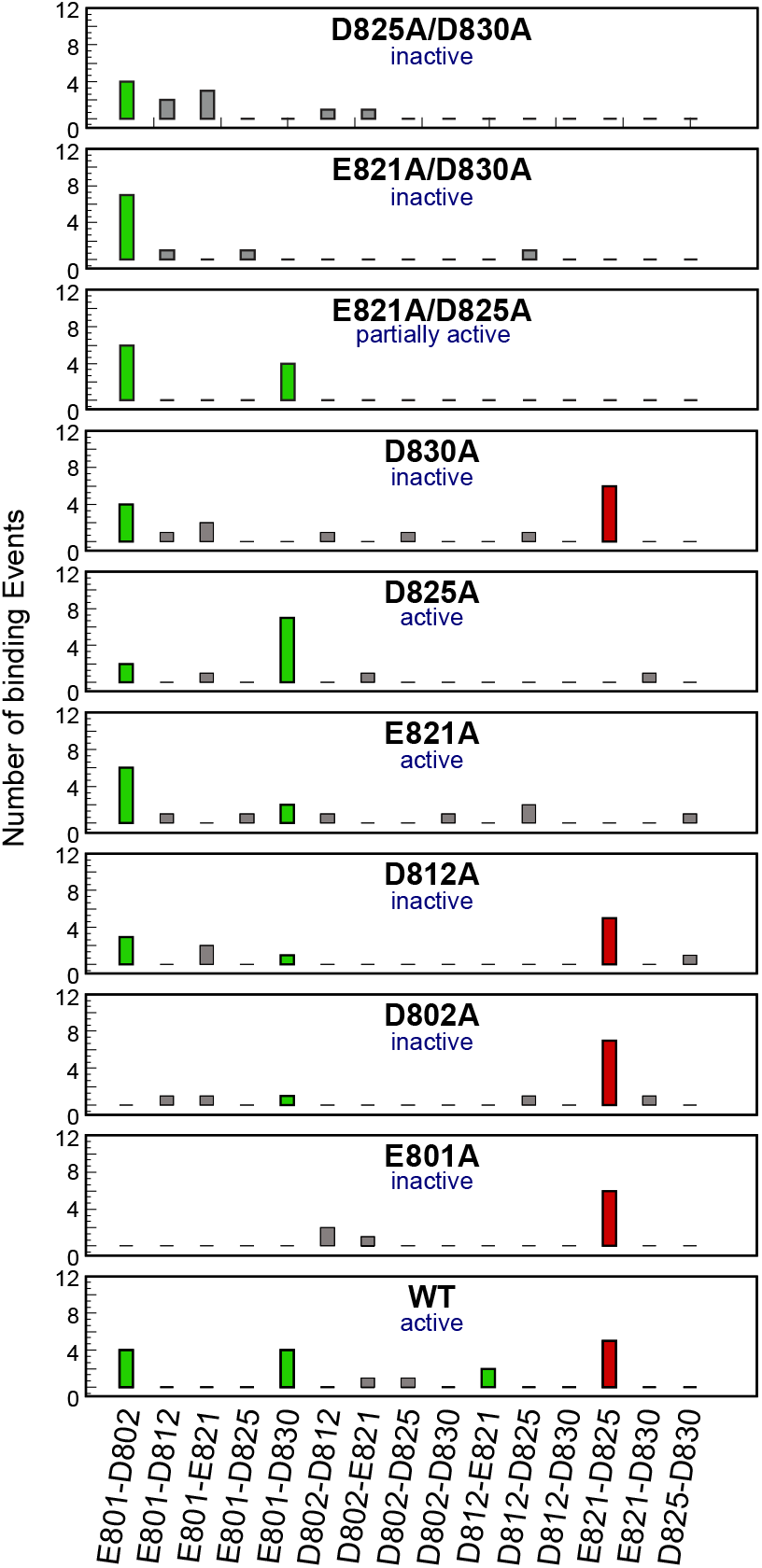
Predicted modes of Ca^2+^ and SARS-CoV fusion peptide binding. Number of Ca binding events to different pairs of anionic residues in SARS-CoV FP in the simulations of the WT and the mutant constructs (labeled in each panel). The experimentally measured phenotypes for each construct are shown (active or inactive). The mode of Ca ion association predicted to be inhibitory for membrane insertion (E821-D825) is shown in red, and the modes of Ca association predicted to facilitate membrane insertion are depicted in green.

From these MD simulations of the SARS-CoV-1 FP constructs, we were able to predict the phenotypes for each construct (active, partially active, or inactive). The mode of calcium ion association predicted to be inhibitory for membrane insertion (E821-D825) is shown in red, and the modes of calcium association predicted to facilitate membrane insertion are depicted in green boxes (**Fig 4**). In those single mutant constructs that were predicted to not have fusion activity (E801A, D802A, D812A, and D830A) we found that the only calcium-coordination mode that persists involves the E821/D825 pair, but not the D812/E821 pair of residues (**Fig 4**). Conversely, in the SARS-CoV-1 FP single mutants that maintained WT-like fusion activity (E821A and D825A), we did not observe the E821/D825 residues to participate in calcium binding. In these function-preserving mutants we identified additional modes of calcium binding that are enhanced in comparison to the WT system: E801/D802 in mutant E821A, or E801/D830 in mutant D825A. For the SARS-CoV-1 FP constructs with multiple mutations, our MD trajectories revealed an overall reduced calcium binding ability (**Fig 4 and SI Fig S3**).

### SARS-CoV-1 FP propensity for membrane insertion is regulated by modes of calcium binding

Overall, the above computational results reveal that the calcium binding patterns of SARS-CoV-1 FP are very similar to those of SARS-CoV-2 FP (21). On this basis, membrane insertion of the SARS-CoV-1 FP could be expected to be enhanced by the modes of calcium binding involving the E801/D802 and D812/E821 pairs of residues, and to be reduced by the ones involving the E821/D825 pair. To test this premise, we carried out MD simulations of FP constructs with two different modes of binding of the Ca2+ ions to the WT SARS-CoV-1 FP associating spontaneously with the lipid membrane (see Methods). In Mode 1, the two calcium ions bind at the E801/D802 and D812/E821 sites; in Mode 2, the calcium ions bind to the E801/D802 and E821/D825 pairs. The membrane interaction of each construct was simulated in 36 independent replicates, each run for ∼0.9-1.0 µs (see Methods).

Analysis of these trajectories revealed that, indeed, the Mode 1 FP extensively penetrated the membrane, while the membrane insertion of the Mode 2 FP was negligible. This can be seen from the plots presented in **Fig 5** comparing frequencies of membrane insertion for each SARS-CoV-1 FP residue in the simulations of Mode 1 and Mode 2 structures.

**Figure 5.**
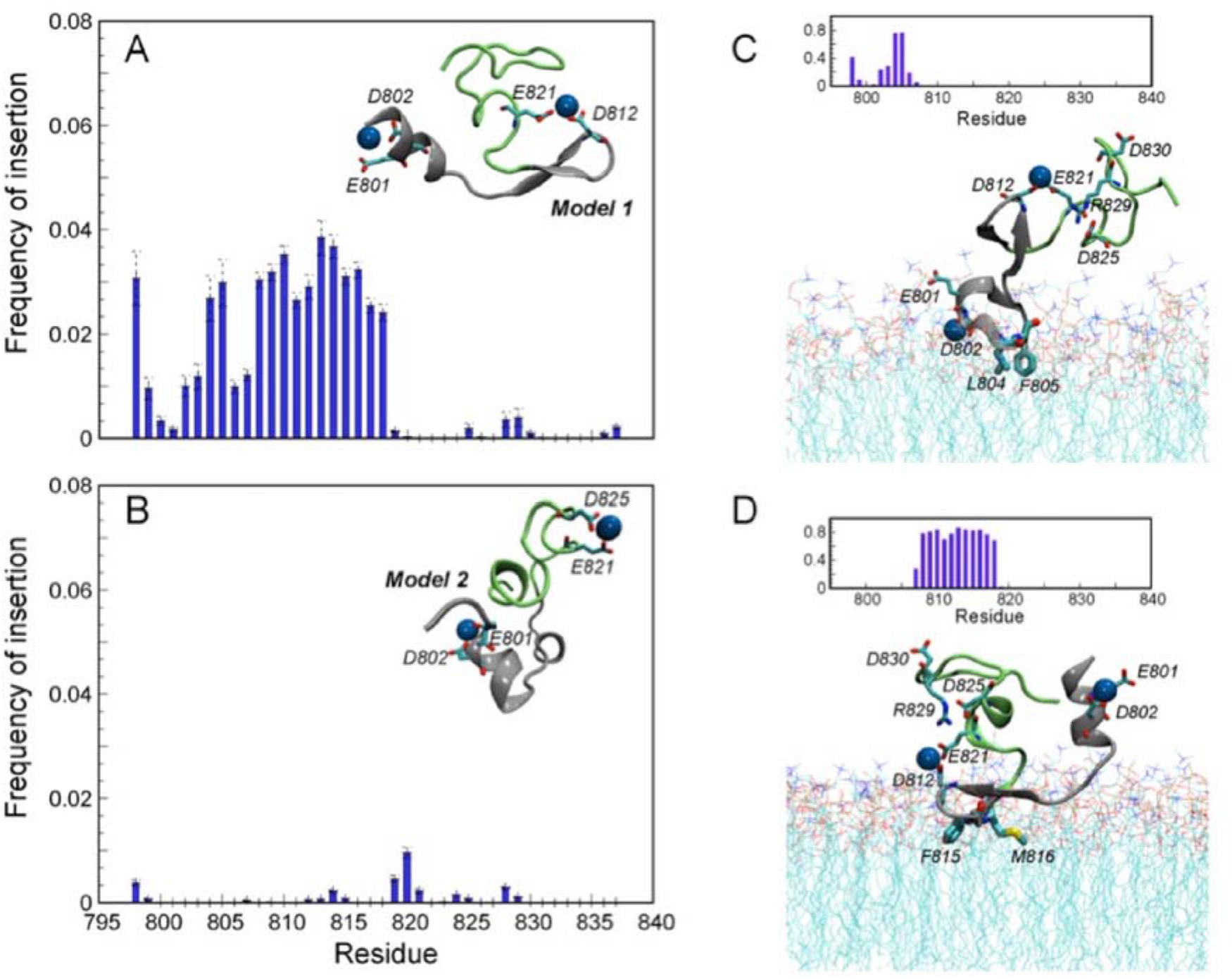
Membrane insertion for FP1 and FP2 domains of SARS-CoV fusion peptide. (A-B) Frequency of membrane insertion for each residue of SARS-CoV FP in the MD simulations of Model 1 (A) and Model 2 (B) constructs, which differ in the mode of Ca^2+^ coordination (see text). The ribbon diagrams in the panels show the corresponding structures. Ca^2+^ coordinating anionic residues are indicated in black text; Ca^2+^ ions are shown as blue spheres. The FP1 and FP2 parts of the fusion peptide are colored in silver and green, respectively. (C-D) The same frequency calculations but done separately for two representative trajectories in Model 1 set in which the observed membrane insertions involved the N-terminal LLF motif (C) and the more centrally located (F815-M816) hydrophobic segment (D). The corresponding ribbon diagrams illustrate the structural features of these two distinct insertion modes highlighting (in black) positions of the anionic residues, the hydrophobic residues penetrating the membrane, and the R829 residue interacting with D825. The same color code applies here as in panels A-B.

The extent of membrane insertion was quantified by monitoring the z-coordinate of the C_α_ atom of each FP residue, with a residue considered to be inserted into the membrane if the z-distance between its C_α_ atom and the second carbon atom in the tail of a POPC lipid (atom C22 in CHARMM36 notation) was <5Å. There is a strong propensity for bilayer insertion of the fusion peptide with Ca^2+^ binding in Mode 1 (**Fig 5A**), compared to the minimal insertion of the FP with Ca^2+^ binding in Mode 2 (**Fig 5B**). A detailed analysis of individual trajectories in the set of Mode 1 binding, revealed two distinct ways in which these SARS-CoV-1 FP constructs penetrate the membrane, similar to our findings for the SARS-CoV-2 FP (21). In one way, the N-terminal segment containing the LLF motif penetrates the bilayer, as shown in **Fig 5C**; in the other, the more centrally located hydrophobic F815-M816 segment enters the bilayer (**Fig 5D**). Interestingly, the two insertion modes appear to alternate which calcium ion binding site is neighboring the inserted portion. Thus, when the LLF is inserted, the calcium ion associated with the neighboring E801/D802 residues is bound to the membrane while the other calcium binding site (D812/E821) is situated away from the membrane surface (**Fig 5C**). In the case of the F815-M816 insertion, the position of the calcium binding loci with respect to the membrane is reversed – the one associated with the D812/E821 pair is membrane-bound, while the E801/D802 pair is located farther from the bilayer (**Fig 5D**). We also note that in both cases, the remaining anionic residues in the peptide (i.e. the ones not engaged with the calcium ions) are either solvent exposed (D830) or engaged in charge-neutralizing interactions with neighboring basic residues (D825/R829) (**Fig 5C and D**). These results support a mechanistic model in which membrane penetration of the SARS-CoV-1 FP is significant only for specific modes of calcium binding to the peptide, i.e., to the E801/D802 and D812/E821 pairs of conserved acidic residues. Moreover, calcium binding at the E821/D825 pair is predicted to inhibit membrane insertion because it leaves charged residues in the vicinity of the hydrophobic conserved LLF unscreened and thus repelled by the negatively charge phospholipid headgroups.

### Analysis of SARS-CoV-1 FP double mutants using syncytia and pseudoparticle transduction assays

We then tested the biological consequences of eliminating pairs of residues from the FP, generating the following double mutants: E821A/D830A, D825A/D830A, and E821A/D825A (**Figure 6A**). As with the single charge-to-alanine S mutants, we validated these mutants by transiently expressing the mutant constructs in HEK-293T cells, biotinylating all cell surface proteins, performing a Steptavidin pulldown, and carrying out spike immunoblots. All three double mutants have comparable cell surface levels of full-length protein compared to WT S, indicating that they are similarly synthesized and trafficked to the plasma membrane (**SI Fig S1B, odd lanes**). In addition, the nearly complete disappearance of the full-length mutant S species and the appearance of the S_1_ and S_2_ subunits following TPCK-Trypsin treatment indicates that these mutants are cleaved similarly to WT S (**SI Fig S1B, even lanes**).

**Figure 6.**
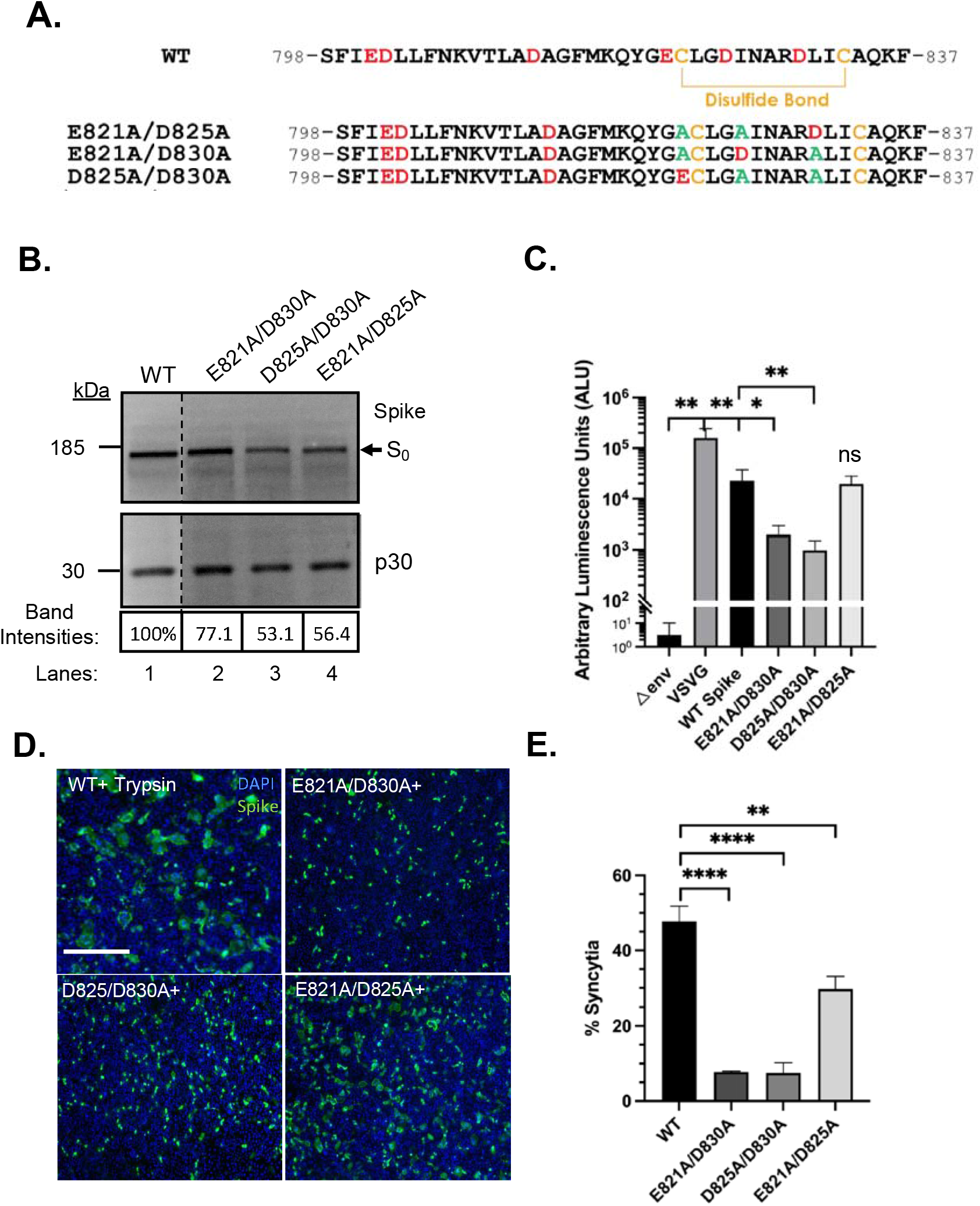
Syncytia formation and pseudoparticle transduction assays of SARS-CoV-1 double mutants. (A) Primary amino acid sequences of the WT and charge-to-alanine SARS-CoV-1 FP double mutants generated (E821A/D830A, D825A/D830A, and E821A/D825A). WT charged residues appear in red, alanine substitutions in green, and cysteine residues in yellow. (B) Representative Spike and MLV p30 immunoblots of pseudoparticles containing WT (lane 1) or double mutant (lanes 2 to 4) SARS-CoV-1 Spike pseudoparticles. S0 indicates the full-length, uncleaved Spike. p30 was used as a loading control. Dashed lines indicate where the WT lanes from the same blots have been reordered for clarity. (C) SARS-CoV-1 pseudoparticle transduction assay in VeroE6 cells with WT and double mutant pseudoparticles. * and ** denote a significance of p ≤ 0.05 or 0.01, respectively, from luminescence values taken from 3 biological replicates. (D) Representative immunofluorescence images of VeroE6 cells transiently expressing WT or the double mutant (E821A/D830A, D825A/D830A, or E821A/D825A) Spike constructs. 24 h following transfection, cells were treated with Trypsin (+ Trypsin or +) for 5 minutes to cleave S proteins and induce syncytia formation. Syncytia were visualized using a SARS-CoV S antibody (green) following cell fixation. Nuclei were stained with DAPI and appear in blue. Images were taken at 20X magnification. Scale bars = 90µm. Images were collected from at least 3 independent experiments. (E) Quantification of percent syncytia across the WT and double mutant conditions. Syncytia were defined as any multinucleated cell with 4 or more nuclei. ** and **** denote significance of p ≤ 0.01 and 0.0001, respectively. n = 3 or 4 biological replicates.

The pseudoparticles containing these double mutants in the SARS-CoV-FP were assayed for incorporation via Spike immunoblotting, which showed that they incorporated their respective S proteins but exhibited variable S protein levels in comparison to the WT pseudoparticles (**Fig 6B**). Transduction assays were then carried out using VeroE6 cells to assess the ability of these pseudoparticles to fuse with and enter host cells. In comparison to WT, the E821A/D830A and D825A/D830A pseudoparticles showed an 11.7-fold and 23.2-fold reduction in cell transduction, respectively (**Fig 6C**).

Given that the D830A single mutation resulted in the greatest reduction in pseudoparticle transduction amongst the single mutants (down 6.8-fold compared to WT pseudoparticles), it is not surprising that the double mutants containing this mutation, in addition to mutations in residues predicted to be non-essential by MDS, remain defective in cell entry. Additionally, these mutant pseudoparticles both had reduced levels of S, which may partially account for their lower cell transduction. In contrast, the E821A/D825A double mutant showed WT-like luminescence values, indicating that mutating both of these residues does not appear to have an effect on pseudoparticle transduction (**Figure 6C**).

We then tested the functionality of these S mutants using the previously described syncytia assay. VeroE6 cells transiently expressing the double mutants E821A/D830A (7.8% syncytia) and D825A/D830A (7.5% syncytia) displayed a pronounced defect in syncytia formation (< 10% syncytia) in comparison to WT S-expressing cells (47.7% syncytia) (**Fig 6D and E**). The E821A/D825A double mutant exhibited a modest decrease in syncytia formation (29.7% syncytia). We observed the same general trends for the double mutants in the pseudoparticle transduction and syncytia assays, namely that the D830A-containing mutants had the greatest detrimental effect, with the E821A/D825A having a smaller effect on syncytia formation but not on pseudoparticle entry. The results from MD simulations for the SARS-CoV-1 FP double mutants studied experimentally (**Fig. 6**) are consistent with these experimental findings. We found the double mutants have an overall reduced calcium binding ability, i.e., mostly a single calcium ion was binding to the peptide at any given time (see details in **SI Fig S3**). The pattern of calcium binding in the functionally deficient double mutants, E821A/D830A and D825A/D830A, was different from that in the partially active E821A/D825A construct (**Fig. 4**). Thus, in the E821A/D825A mutant, calcium preferentially bound to the E801/D830 pair, which is a preferred mode of binding in the WT as well, while the other two constructs are missing the D830 residue which was shown to be important for function.

## Discussion

Early structure-function studies of the SARS-CoV-1 FP pointed to peptide regions – especially the largely conserved LLF motif and the residues flanked by cysteines (C822-C833) – as important features for membrane fusion activity (22). In this work, a charge reversal point mutation (D830L) and a deletion of several amino acids including this residue (Δ823-832) were found to have pronounced effects on membrane fusion and pseudoparticle transduction (22); however, these studies were performed before the importance of calcium during viral entry was known and hence their mechanistic importance, described here, was missed. More recent biophysical characterization of the FP has found that mutating residue D812 in the FP1 region has the greatest effect on membrane ordering, calcium-specific FP binding, and calcium-induced FP structural changes (23). Residue D830 in the FP2 region (labeled 831 in the citation) was also found to be important, causing loss of membrane ordering and calcium sensitivity when mutated, as were residues 802 and 801, which they proposed together form a calcium binding site (23).

In this systematic study of the roles of the negatively charged residues in the SARS-CoV-1 FP, we first confirmed their importance for pseudoparticle transduction, a proxy we use for viral infectivity. After validating the mutant S proteins (**SI Fig S1**) and confirming their incorporation into pseudoparticles (**Fig 2A**), we found that SARS-CoV-1 pseudotyped particles containing either the Spike D802A or D830A mutation had decreased transduction into VeroE6 cells (**Fig 2B**). Conversely, D812A and D825A S-containing pseudoparticles were as able to enter cells as well as their WT counterparts. An interesting finding from this set of experiments is that the E821A mutation enhanced transduction, suggesting that it is advantageous during viral entry. The test of fusion competency of the single FP mutants showed that VeroE6 cells expressing the D802A or D830A mutants exhibited a pronounced cell-cell fusion defect, while the D812A mutant showed a more moderate decrease. The D821A and D825A mutants formed syncytia similar to WT S-expressing cells (**Fig 3**). In summary, our biological data from our single mutants confirm that residues D802 and D830 are essential in a non-redundant manner to the function of the SARS-CoV-1 fusion peptide and that loss of transduction ability in these pseudoparticle mutants is likely due to a defect at the level of membrane fusion.

We then conducted MD simulations of the WT and mutant constructs (summarized in **Fig. 4**) to identify the preferred modes of SARS-CoV-1 FP-Ca^2+^binding and interrogate which negatively charged residues, when bond to calcium, promote membrane insertion (**Figs. S2** and **S3** in **SI**). We anticipated that FP residues D802 and D830 bound to calcium ion would position the FP favorably for membrane insertion, given that their genetic ablation resulted in loss of S protein fusion competency and decreased pseudoparticle transduction. Indeed, in our simulations, FP residues D802 and D830 were predicted to be part of a pairwise interaction with calcium ions that promotes membrane insertion, Furthermore, single mutations in either of these residues were predicted to inactivate the FP. More broadly, the energetically favorable modes of Ca^2+^-FP binding identified to promote membrane insertion (**Fig. 5**) all spatially organize the FP to encourage its insertion by (i)-charge screening, and (ii) -stabilizing the peptide conformation for penetration of the hydrophobic LLF motif beyond the phospholipid headgroups. Our results from probing the effect of pairwise mutations of the negatively charged residues in the SARS-CoV-1 peptide support the roles of specific pairs of Ca^2+^ binding residues.

The results of MD simulations of the double mutant constructs (**Figs 4**) are fully compatible with the expectations from the experimentally validated predictions for the single mutant constructs (**Fig. 3 and Fig. 4**) and the experimental results shown in **Fig. 6**. The depth of insertion measured in the simulated FP-inserted systems, and the determinant role of the calcium ion in facilitating membrane insertion are identical to the findings for the SARS-CoV-2 peptide (21) and are in agreement with the experimental measurements of SARS-CoV-1 FP membrane insertion by Lai et al (17, 18, 32). That calcium interacts with the same highly conserved, negatively charged residues in the FPs of SARS-CoV-1 and SARS-CoV-2 is not surprising, given the high sequence similarity between the two FPs.

The results from the combined experimental and computational probing of SARS-CoV-1 FP-membrane interactions reveal that the propensity of SARS-CoV-1 FP membrane insertion is determined by specific modes of calcium binding. The calcium binding mode that enables the most sustained membrane penetration involves the initial association of the peptide with the bilayer through a calcium binding site located near the N-terminus of the peptide. This site (E801/D802) is made accessible following enzymatic cleavage and is inserted into the membrane in conjunction with the insertion of the juxtaposed hydrophobic segment (the LLF motif). In this process, all anionic residues of the peptide are engaged with calcium ions, or with neighboring basic residues, unless they remain solvent exposed away from the membrane. Based on the preferred modes of calcium-loaded FP interaction with the membrane generated from our MD simulations, we conclude that the binding of calcium following S protein cleavage at the S2’ site creates an energetically feasible membrane insertion process. This process consists of the following three steps: (1) steering of the peptide towards the bilayer surface by shielding calcium interactions with the phospholipid headgroups; (2) enabling the conserved N-terminal LLF motif of the FP to reach the hydrophobic layer of the membrane by screening the negative charges of the neighboring residues involved in Ca binding; and (3) shielding of FP anionic residues in the preferred inserting-enabling conformation of the peptide to prevents negative charge from interfering with membrane insertion (**Fig 5**). These three components became evident from the MD simulations that identified the most likely modes of Ca binding, the preferred conformations adopted by Ca loaded FPs, and the effect of the mutations on these properties and insertion probabilities.

Not only do the molecular mechanisms presented here for the SARS-CoV-1 FP bear a very high similarity to results from computational studies of the SARS-CoV-2 FP (21), but in both the SARS-CoV-2 FP simulations (33) and here, we observed a Ca^2+^ binding mode that inhibits stable membrane insertion. In this mode, calcium association with the E821/D825 pair (corresponding to the D839/D843 in SARS-CoV-2 FP) positions the LLF motif near the negatively charged D812 residue at the membrane surface (**Fig 5B**). This eliminates the favorable effect of the negative charge screening by the calcium, so that the peptide bilayer encounters are transient and do not lead to penetration by the LLF motif. This mechanistic model, which highlights the role of the positioning of the D812 residue in specific calcium binding modes, is supported by our findings for the SARS-CoV-1 S mutants, as it was previously found to be an impactful mutation in the SARS-CoV-1 FP (20). Furthermore, we directly tested the biological consequences of generating the single and double E821A/D825A mutants, observing that although the E821A mutant pseudoparticles showed an increase in transduction, the single D825A single mutant and E821A/D825A double mutant did not (**Fig 6E**).

Indeed, data in **Figs 4** and **5** show that mutations that create a high propensity for the calcium binding mode involving the E821/D825 pair, stabilize peptide conformations that are non-productive for membrane penetration and are found here to inactivate the FP. Current studies on SARS-CoV-2 S protein classified the disordered portion of the FP in the FP2 domain as the FPPR (fusion peptide proximal region), which includes the charged residues D839/D843/D848 or E821/D825/D830 equivalent in SARS-CoV-1 FP (21, 34). This FPPR region was determined to be important, as it binds to the RBD through the CTD1, and maintains the closed pre-fusion S trimer(34). This is due to the tight packing around the disulfide bond reinforced by a bond between K835 and D848 (K817 and D830 in SARS-CoV-1 FP). The equivalent residue in SARS-CoV-1 FP, based on biological studies, is a critical residue (34). Mutating D830 removes the binding partner of K817, thereby loosening the ‘knot’ of the structured FPPR in maintaining the closed form of the trimeric S protein (34). Without the reinforcements, we may lose the optimal structural conformation of the S protein upon receptor binding to expose the S2’ cleavage site to promote successful proteolytic cleavage.

A recent study from Koppisetti et al asserts that a calcium ion may be binding to D843/D849 in SARS-CoV-2 FP (corresponding to D825/D830 in SARS-CoV-1 FP); however, as shown from the simulations done in this study and our previous report (21), there is a high likelihood that the second calcium ion will bind to D812/D821 (corresponding to the D830/D839 in SARS-CoV-2 FP) for the maximal membrane penetration (21, 35). We note that the model of membrane insertion proposed by Koppisetti et al (35) most aligns with our proposed CoV FP interaction with the bilayer (**Fig 5**), but that NMR studies regarding the membrane interaction with the fusion peptide use bicelles (35) and micelles (36) which are not faithful models for the cell membrane bilayer.

More broadly, the E801 and D830 residues are conserved within the CoV family and the D812 residue is conserved within betacoronaviruses (17, 20). The implication is that calcium interactions are a conserved mechanism that serves to better position the FP for membrane insertion. Different coronaviruses exhibit different requirements for calcium; MERS-CoV binds to one calcium ion in its FP1 domain (18), thus, it is important to investigate the role of calcium and FP interactions across the CoV family. Recent work using a FRET-assay has shown that calcium interactions with the fusion peptide may drive the pre- to post-fusion conformational changes in SARS CoV-2 spike (31). The authors of this study showed that alpha and beta coronaviruses display a high sensitivity to calcium; the fusion activity of spike drops precipitously as calcium levels rise or decrease past predicted late endosomal concentrations. The conservation of calcium-binding residues in the FP of many coronaviruses suggests that the CoV fusion mechanisms are essential for function and can serve as potential targets for broad-spectrum antiviral drugs (37–39). Repurposing FDA-approved calcium channel blocking (CCB) drugs to inhibit CoV entry, particularly for SARS-CoV-2, is one option worth exploring. Recent studies have shown that the CCB felodipine, among others, is a potential therapeutic candidate to inhibit SARS-CoV-2 entry (40). CCBs can target conserved viral functions, providing a rapid solution to address new and future SARS-CoV-2 variants. It will be important to identify the mechanisms of CCBs CoV inhibition, as they may directly inhibit a viral target or indirectly inhibit viral entry by affecting host cell processes. Interestingly, SARS-CoV-2 variants have arisen as part of Clade 20A that contain a D839G or D839Y mutation (E821 equivalent in SARS-CoV-1), with this mutation predicted to affect FP-Calcium interactions (41–43). To date, it is not known if there is any selective advantage to the virus conferred by this mutation or whether the emergence of these variants simply represents a founder effect. Future studies will continue to illustrate the role of the spike FP in controlling both emerging and existing coronaviruses.

## Conclusion

In the field, how calcium binding specifically impacts the dynamics of coronavirus fusion peptide dynamics during host membrane insertion was lacking. We have taken a two-pronged approach to directly investigate this aspect of SARS-CoV-1 entry, using biological studies in combination with molecular dynamics simulations to identify which anionic FP residues interact with calcium ions. We have also demonstrated how specific pairwise interactions likely impact FP function, and ultimately, virus entry. Interestingly, we have found that multiple binding modes of FP-Ca^2+^ exist, with one mode promoting FP membrane insertion and another inhibiting membrane insertion. Our predictive studies are largely in agreement with the findings of our biological data, demonstrating the power of this cross-disciplinary approach to get at viral entry mechanisms.

## Materials and Methods

### Cells, plasmids, and reagents

Human embryonic kidney 293 (HEK293T) and African green monkey kidney epithelial (VeroE6) cells were obtained from the American Type Culture Collection (ATCC, Manassas, VA). Both cell lines were grown in complete DMEM, composed of Dulbecco’s Modified Eagle medium (DMEM, CellGro), supplemented with 10% HyClone FetalClone II (GE) and 10 mM HEPES (CellGro), and grown at 37°C with 5% CO_2_. The plasmids used for generating pseudoparticles were the pCMV-MLV gag-pol murine leukemia virus (MLV) packaging construct, the pTG-Luc transfer vector encoding the luciferase reporter gene, and the pCAGGS-VSVG plasmids. They were generously provided by Jean Dubuisson (Lille Pasteur Institute, Lille, France) and co-transfected as previously described (26). The plasmid encoding the C9-tagged SARS-CoV-1 spike protein (pcDNA3.1-C9-SARS-CoV-1 S) was provided by Dr. Michael Farzan from the New England Primate Research Center. Recombinant L-1-tosylamide-2-phenylethyl chloromethyl ketone (TPCK)-treated trypsin was obtained from Sigma.

### Site-directed mutagenesis

Site-directed mutagenesis was performed on the SARS-CoV-1 spike encoding plasmid, pcDNA3.1-SARS-CoV-1-S, via the QuikChange Lightning site-directed mutagenesis kit (Aligent). PCRs and transformations were performed based on the manufacturer’s recommendations. The primers used to generate the SARS-CoV-1 S mutants can be found in the supplementary information (**SI, Table S1**). Mutations were confirmed via Sanger Sequencing at the Cornell University Life Sciences Core Laboratories Center.

### Cell Surface Biotinylation Assay

HEK293T cells were seeded in poly-D-lysine coated 6-well dishes. 24 hours later, cells were transfected with 1.5 ug of WT or mutant SARS-CoV-1 S plasmid using polyethylenimine (PEI) (Thermo Fisher Scientific). Cells were transfected for 24 hours, washed once with 1X DPBS, and then left untreated or treated with 1 ug/mL of TPCK-trypsin in DPBS for 5 minutes at 37°C. Cell surface proteins were then biotinylated by incubating rinsed cells in a biotin-containing buffer (250 ug/mL in PBS) for 20 minutes. The biotin buffer was then replaced with a quenching solution (50mM glycine in DPBS) and incubated for 30 minutes. Cells were then lysed in ice for 15 minutes using a lysis buffer (0.1% Triton in 1x TBS) with protease inhibitors (cOmplete Protease Inhibitor Cocktail). Lysates were collected and spun at 13,000 rpm for 10 minutes to pellet insoluble material. The lysates were then incubated overnight at 4° C with equilibrated steptavidin beads to retrieve the biotinylated spike proteins. The captured spike proteins were eluted by boiling the beads in 1x LDS sample buffer in the presence of DTT (NuPAGE) for 10 minutes. Proteins were resolved on 4-12% gradient Bis-Tris gels (NuPAGE) and transferred to PVDF membranes. The SARS-CoV-1 S protein was detected using the SARS-CoV-1 S rabbit polyclonal primary antibody (NR-4569, BEI resources) and Alexa Fluor 488-labeled goat anti-rabbit secondary antibody (Invitrogen). The resulting protein bands were visualized using a Chemidoc system with Image Lab image capture software (BioRad).

### Cell-cell fusion assay

VeroE6 cells were seeded onto glass slides in a 24-well chamber slides (Millipore). After 24 hours, cells were transfected using Lipofectamine according to the manufacturer instructions. Following 18 hours of transfection, cells were washed with 1X DPBS and Trysinized with 2ug/mL TPCK-trypsin for 5 minutes at 37°C to. Cells were then fixed with 4% paraformaldehyde (PFA) (ThermoFisher) for 15 minutes and washed three times with DPBS. To permeabilize the cells, a 0.1% Triton X-100 in DPBS solution was added to each well and cells were incubated for 5 minutes. After three washes with DPBS, the cells were blocked with 5% normal goat serum for 30 minutes. Cells were again washed and then labeled with SARS-CoV-1 S rabbit polyclonal antibody (NR-4569, BEI resources), followed by labeling with Alexa Fluor 488-labeled goat anti-rabbit secondary antibody (Invitrogen). The cell nuclei were labeled with a DAPI stain present in the mounting media (DAPI Fluoromount-G, Southern Biotech). Images were acquired using an upright microscope (Echo Revolve) with a 10x objective. To quantify the number of nuclei per syncytia, three randomly selected fields were acquired and spike-expressing cells and syncytia were manually counted. The total number of syncytia we counted and expressed as a percentage of the total fluorescent “particles” that were counted in the image. The data was graphed and statistics were carried out using Microsoft Excel and GraphPad Prism 7.

### SARS-CoV-1 pseudoparticle infectivity assay

SARS-CoV-1 pseudoparticles were produced as previously described (18, 26). Briefly, to generate pseudoparticles 3.5 x10^5^ HEK293T cells were seeded in 6-well plates. Pseudoparticles were prepared by transfecting HEK293T cells with 600 µg of their respective SARS WT or mutant S plasmids, 800 µg of pTG-Luc plasmid, and 600 µg of pCMV-MLV gag-pol plasmid using polyethylenimine (PEI) as the transfection reagent. The cell supernatant was harvested 48 hours post-transfection, centrifuged at 1200 rpm for 7 minutes to separate the pseudoparticles from residual cellular debris, and filtered through a 0.45 µm syringe filter. The pseudoparticles were stored at -80°C for one freeze-thaw cycle. To perform the pseudoparticle infectivity assay, 5×10^5^ VeroE6 cells were seeded into 24-well plates and infected with 200uL of the pseudoparticle suspension for 72 hrs at 37°C. Cells were lysed for 10 minutes at room temperature using the Luciferase cell lysis reagent (Promega). Luminescence readings were performed using a Glomax 20/20 Luminometer system (Promega). Each experiment contained a minimum of three technical replicates and experiments were repeated at least three times. Data analysis was performed using GraphPad Prism 7.

### Modeling of the SARS-CoV-1 fusion peptide

The structural model of the SARS-CoV-1 S protein fusion peptide is based on the SARS-CoV-1 S prefusion structure from the Protein Data Bank, PDB 5XLR. The sequence of the SARS-CoV-1 S protein (GenBank accession no. AAT74874.1) was aligned to the PDB 5XLR SARS-CoV-1 structure sequence using Geneious software (v.2020.1.1). Structural models of the SARS-CoV-1 S monomer were generated using the Modeller comparative modeling tool (**v.9.23**) within the Chimera software (**v.1.13**; University of California). Images were created using Adobe Illustrator CC (**v.24.03**).

### Molecular dynamics (MD) simulations of the SARS-CoV-1 FP in water

For all the atomistic MD simulations, the SARS-CoV-1 FP segment was capped with neutral N- and C-termini (ACE and CT3, respectively, in the CHARMM force-field nomenclature). Protonation states of all the titratable residues were predicted at pH 7 using Propka 3.1 software (44).

For the simulations in water, one copy of the peptide (wild-type or a mutant) was embedded in a rectangular solution box and ionized using VMD tools (“Add Solvation Box” and “Add Ions”, respectively) (45). The box of dimensions ∼90 Å x 80 Å x 82 Å included a Na^+^Cl^-^ ionic solution as well as 2 Calcium ions, and ∼18000 water molecules. The total number of atoms in the system was ∼54,540.

The system was equilibrated with NAMD version 2.13 (46) following a multi-step protocol during which the backbone atoms of the SARS-CoV-1 FP as well as Calcium ions in the solution were first harmonically constrained and subsequently gradually released in four steps (totaling ∼3ns), changing the restrain force constants k_F_ from 1, to 0.5, to 0.1 kcal/ (mol Å^2^), and 0 kcal/ (mol Å^2^). These simulations implemented all option for rigidbonds, 1fs (for k_F_ 1, 0.5, and 0.1 kcal/ (mol Å^2^)) or 2fs (for k_F_ of 0) integration time-step, PME for electrostatics interactions (47), and were carried out in NPT ensemble under isotropic pressure coupling conditions, at a temperature of 310 K. The Nose-Hoover Langevin piston algorithm (48) was used to control the target P = 1 atm pressure with the “LangevinPistonPeriod” set to 200 fs and “LangevinPistonDecay” set to 50 fs. The van der Waals interactions were calculated applying a cutoff distance of 12 Å and switching the potential from 10 Å.

After this initial equilibration phase, the velocities of all atoms in the system were reset and ensemble MD runs were initiated with OpenMM version 7.4 (49) during which the system was simulated in 18 independent replicates, each for 640ns (i.e., cumulative time of ∼11.5 µs for each FP construct). These runs implemented PME for electrostatic interactions and were performed at 310K temperature under NVT ensemble. In addition, 4fs time-step was used, with hydrogen mass repartitioning and with “friction” parameter set to 1.0/picosecond. Additional parameters for these runs included: “EwaldErrorTolerance” 0.0005, “rigidwater” True, and “ConstraintTolerance” 0.000001. The van der Waals interactions were calculated applying a cutoff distance of 12 Å and switching the potential from 10 Å.

### MD simulations of SARS-CoV-1 FP interactions with lipid membranes

Interactions of selected two models of the Calcium-bound WT SARS-CoV-1 FP with lipid membranes were investigated with atomistic MD simulations. These runs were initiated by placing each of the models in the proximity of a bilayer composed of 3:1:1 POPC/POPG/Cholesterol that had been pre-equilibrated for 25ns as described previously (21).

After the FP-membrane complexes were embedded in a solution box (containing 150 mM Na^+^Cl^-^ salt concentration), each system was equilibrated with NAMD version 2.13 following the same multi-step protocol described above during which the backbone atoms of the FP as well as the Calcium ions were first harmonically constrained and subsequently gradually released in four steps. After this phase, the velocities of all atoms of the system were reset, and ensemble MD runs were initiated with OpenMM version 7.4. Each system was simulated in 18 independent replicates, each ran for ∼ 1 µs (i.e., cumulative time of ∼18 µs for each FP-membrane complex). These runs implemented PME for electrostatic interactions and were performed at 298K temperature under NPT ensemble using semi-isotropic pressure coupling, with 4fs time-steps, using hydrogen mass repartitioning and with “friction” parameter set to 1.0/picosecond. Additional parameters for these runs included: “EwaldErrorTolerance” 0.0005, “rigidwater” True, and “ConstraintTolerance” 0.000001. The van der Waals interactions were calculated applying a cutoff distance of 12 Å and switching the potential from 10 Å. For all simulations we used the latest CHARMM36 force-field for proteins and lipids (50), as well as the recently revised CHARMM36 force-field for ions which includes non-bonded fix (NBFIX) parameters for Na^+^ and Calcium ions (51).

### Supporting Information

**Figure S1** shows representative Spike immunoblots results of our Biotinylation Assays, which we carried out with the wild-type and mutant SARS-CoV-1 constructs used in this study.

**Figure S2** shows the predicted models of calcium ions binding to the wild-type and single charge-to-alanine mutants generated from the MD simulations.

**Figure S3** shows the predicted models of calcium ions binding to the wild-type and double charge-to-alanine mutants generated from the MD simulations.

**Table 1** is a list of the primer sets used to generate the SARS-CoV-1 single and double mutants.

### Corresponding Author Information

Susan Daniel, Ph.D., William C. Hooey Director & Fred H. Rhodes Professor of Chemical Engineering, Robert Frederick Smith School of Chemical and Biomolecular Engineering Cornell University | 124 Olin Hall, Ithaca, NY 14853 USA, phone: (607)255-4675

### Present/Current Author Addresses

Javier A. Jaimes, Ph.D.

The University of Massachusetts Medical School, Program in Molecular Medicine, 373 Plantation Street, Biotech 2 Suite 319 Worcester MA 01605 USA, phone: (508)856-6891

### Author Contributions

Conceived and designed the experiments: MKB JDC GK MRS TT JAJ HW GRW SD. Performed the experiments: JDC GK* MKB MRS TT. Analyzed the data: JDC GK MKB MRS TT JAJ HW GRW SD. Wrote and edited the paper: JDC MKB GK MRS TT HW GRW SD. Acquired funding: GRW SD HW. Computational Resources: HW GK.

## Acknowledgements

We would like to thank members of the Daniel, Whittaker, and Abbott groups as well as the Weinstein group at Weill Cornell for helpful discussions. This work was supported by The National Institute of Health research grant R01AI35270, National Science Foundation RAPID grant 2027070 and Fast Grant from the Mercatus Center at George Mason University. TT is supported by the National Science Foundation Graduate Research Fellowship Program under Grant No. DGE-1650441 and the Samuel C. Fleming Family Graduate Fellowship. H.W. and G.K. gratefully acknowledge support from the 1923 Fund, and the access to computational resources awarded through the COVID-19 High Performance Computing Consortium at the Center for Computational Innovations (CCI) at the Rensselaer Polytechnic Institute and Accounts BIP225 and BIP109 at the Oak Ridge Leadership Computing Facility, which is a DOE Office of Science User Facility supported under Contract DE-AC05-00OR22725.

## Abbreviations Used

Fusion Peptide (FP)

Spike (S)

Molecular Dynamics Simulations (MDS)

Coronaviruses (CoVs)

---

**For Table of Contents Only**

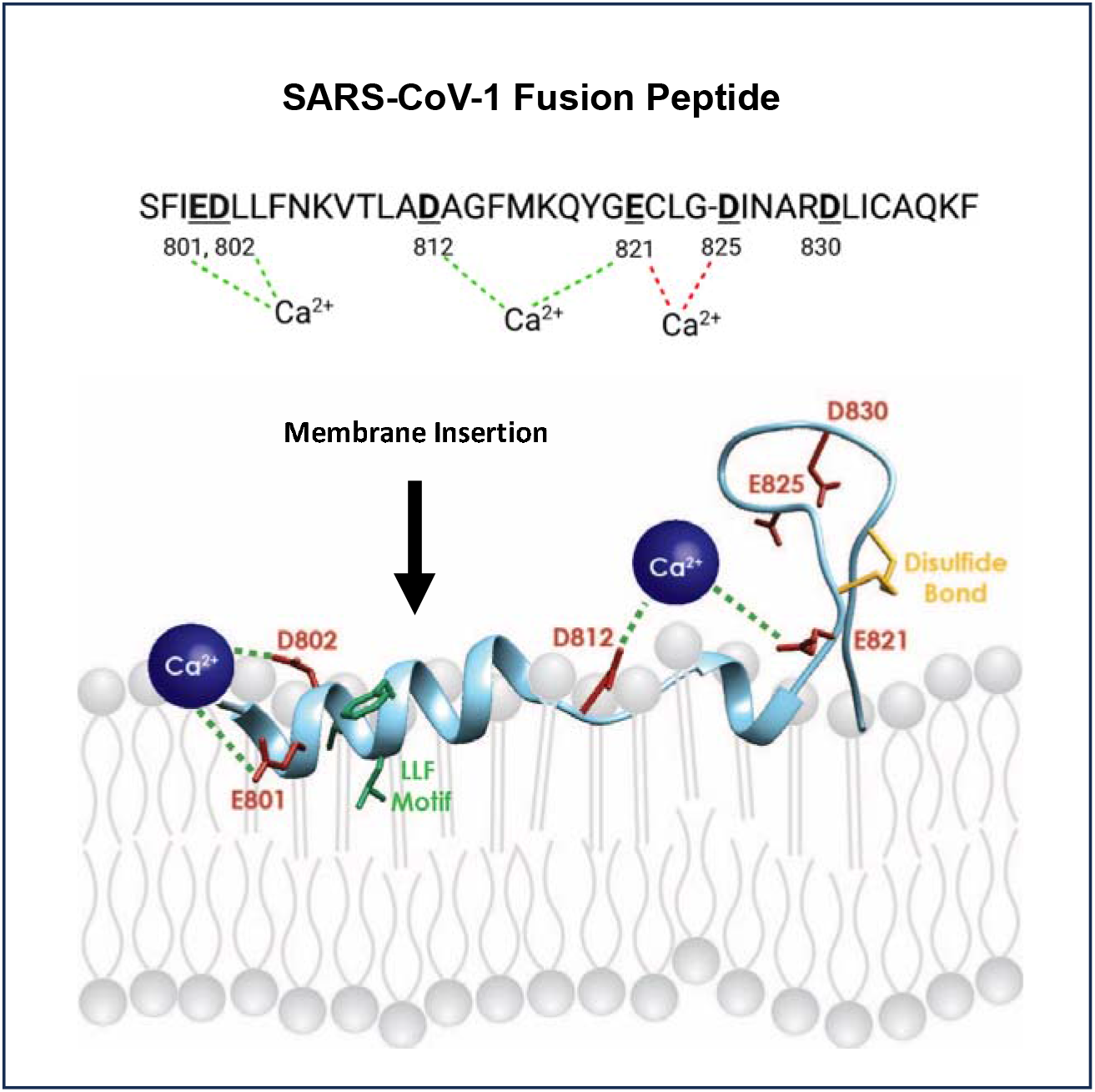

## Notes

### Competing Interest Statement

The authors have declared no competing interest.

### Summary of Updates

While sequencing the plasmids used in this study we discovered that the the labeled D802A and D812A single SARS CoV-1 FP mutants were incorrect, with the wrong residues mutated in each construct (D802A was D812A and D812A was double mutant E801A/D802A). We have corrected the plasmids and rerun all biological assays with the actual D802A and D812 FP mutants. The results of the biological assays have changed with regards to the two corrected mutants. For the pseudoparticle studies, the D802A mutant exhibited a more moderate decrease in transduction in comparison to the previous incorrect mutant. The correct D812A mutant pseudotyped particles were able to transduce VeroE6 cells comparable to WT pseudoparticles, unlike the complete loss of transduction seen with the previous incorrect mutant. Unrelated to the corrected mutants, we saw a significant increase in E821A pseudoparticle transduction, likely due to the increased biological and technical replicates we've completed to repeat these experiments with the full sets of mutants. This differs from our earlier work, in which we saw no difference in pseudoparticle transduction of VeroE6 cells between the WT and mutant E821A conditions. The syncytia assay results have also changed with respect to the corrected mutants. The correct D802A mutant has a more severe defect in syncytia formation in VeroE6 cells, whereas the incorrect previous mutant formed syncytia like WT Spike-expressing cells. The correct D812A mutant has a more moderate decrease in syncytia formation compared to the complete loss in syncytia formation with the incorrect mutant. Upon repeating the transduction assay with additional batches of pseudoparticles, we could not consistently get an increase in transduction with the E821/D825 double mutant, as we previously reported. Instead we find this mutant to transduce VeroE6 cells similarly to WT-containing pseudoparticles. We have updated the pseudoparticle data to reflect this change. Lastly, we have removed the calcium depletion studies because we felt they did not add any new findings to the field, despite being an important validation step.

