## Supplemental Information for "A mechanistic understanding of the modes of Ca ion binding to the SARS-CoV-1 fusion peptide and their role in the dynamics of host membrane penetration"

**Supplemental Information - Figure 1**

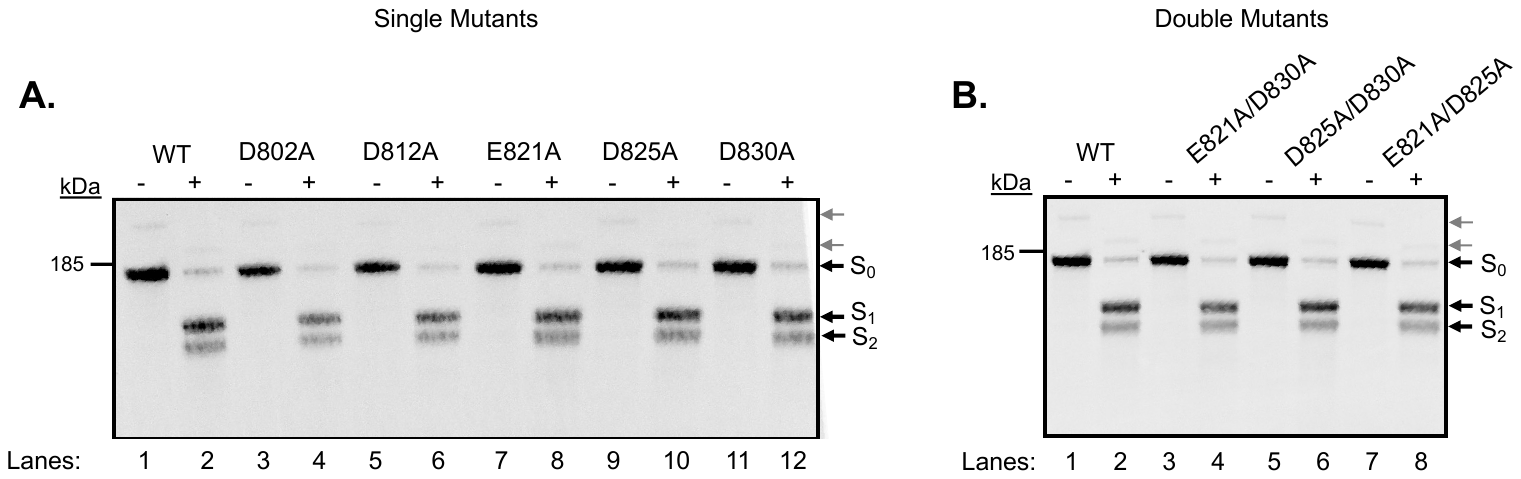

**SI Figure S1.** Cell surface biotinylation assay of SARS-CoV-1 wild-type and charge-to-alanine spike mutants. (A and B) Representative Spike immunoblots of biotinylated WT and single (left) or double (right) mutant S proteins transiently expressed in HEK293T cells, then left untreated (-) or treated (*+*) with 1 ug/mL Trypsin for 10 minutes. S_0_ indicates the full-length, uncleaved spike protein. S_1_ and S_2_ indicate the cleaved subunits of spike following trypsinization. Gray arrows indicate faint higher molecular weight bands observed. n = 3 biological replicates.

**Supplemental Information - Figure 2**

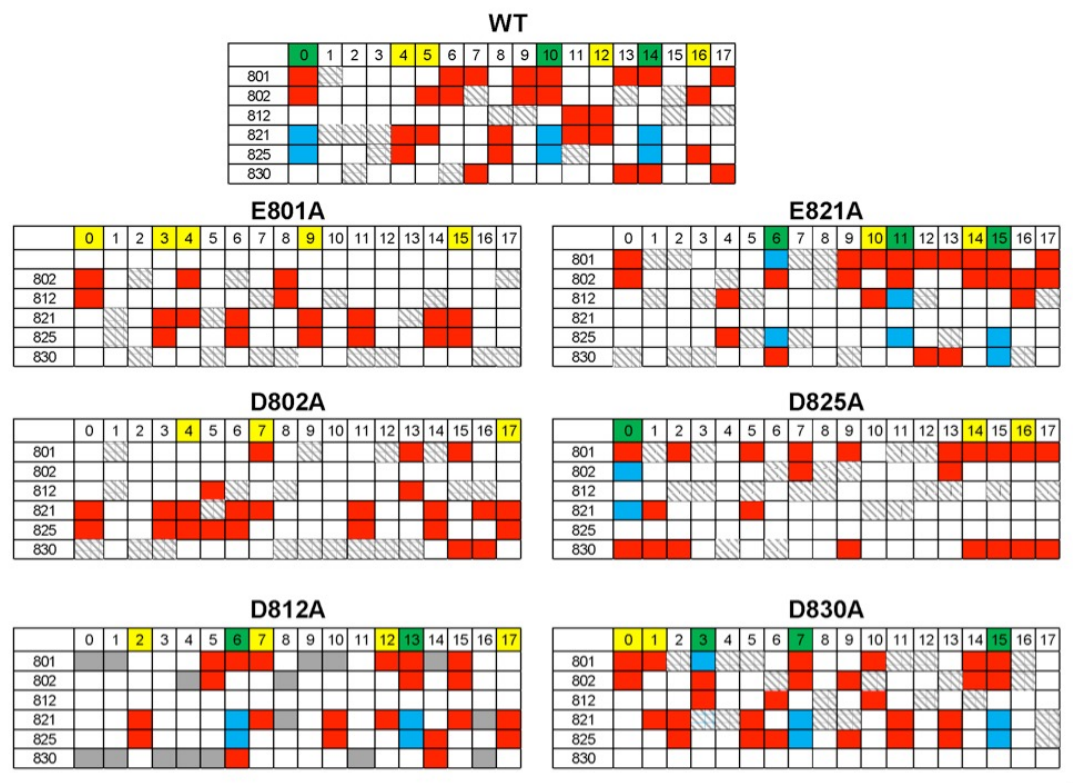

**SI Figure S2.** Models of Ca^2+^ binding to the WT and single mutant SARS-CoV FPs. The tables show, for each construct, acidic residues implicated in Ca^2+^ binding in 18 independent atomistic MD simulations (each 640ns in length). In a particular trajectory, residue pairs simultaneously engaging with the bound Ca^2+^ are denoted by *red* or *blue* rectangles, whereas instances of Ca^2+^ ion associating with a single acidic residue is depicted with *grey-striped* rectangle. The trajectories in which simultaneous binding of two Ca^2+^ ions to different pairs of residues were observed are highlighted in *green*. The simulations in which a single Ca^2+^ ion was bound to a pair of acidic residues are shown in *yellow*.

**Supplemental Information - Figure 3**

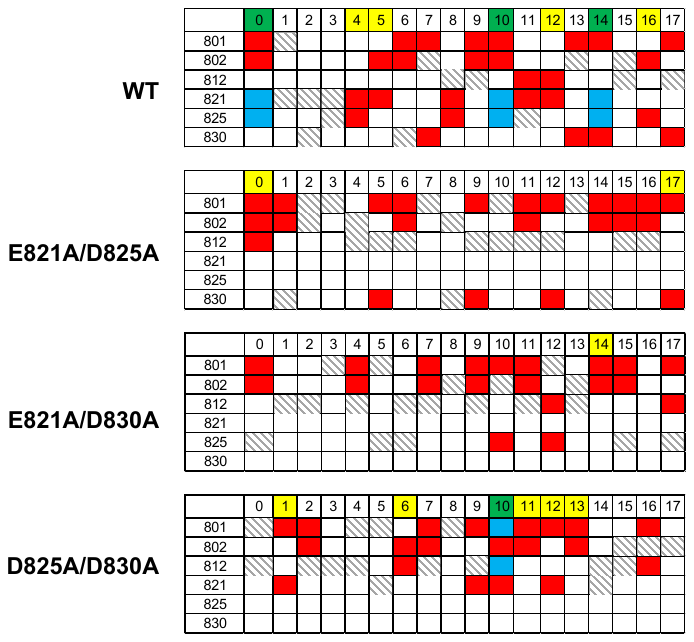

**SI Fig S3.** Models of Ca2+ binding to the WT and multiple mutant SARS-CoV FPs. The tables show, for each construct, acidic residues implicated in Ca2+ binding in 18 independent atomistic MD simulations (each 640ns in length). In a particular trajectory, residue pairs simultaneously engaging with the bound Ca2+ are denoted by *red* or *blue* rectangles, whereas instances of Ca2+ ion associating with a single acidic residue is depicted with *grey-striped* rectangle. The trajectories in which simultaneous binding of two Ca2+ ions to different pairs of residues were observed are highlighted in *green*. The simulations in which a single Ca2+ ion was bound to a pair of acidic residues are shown in *yellow*.

**Supplemental Information - Table 1**

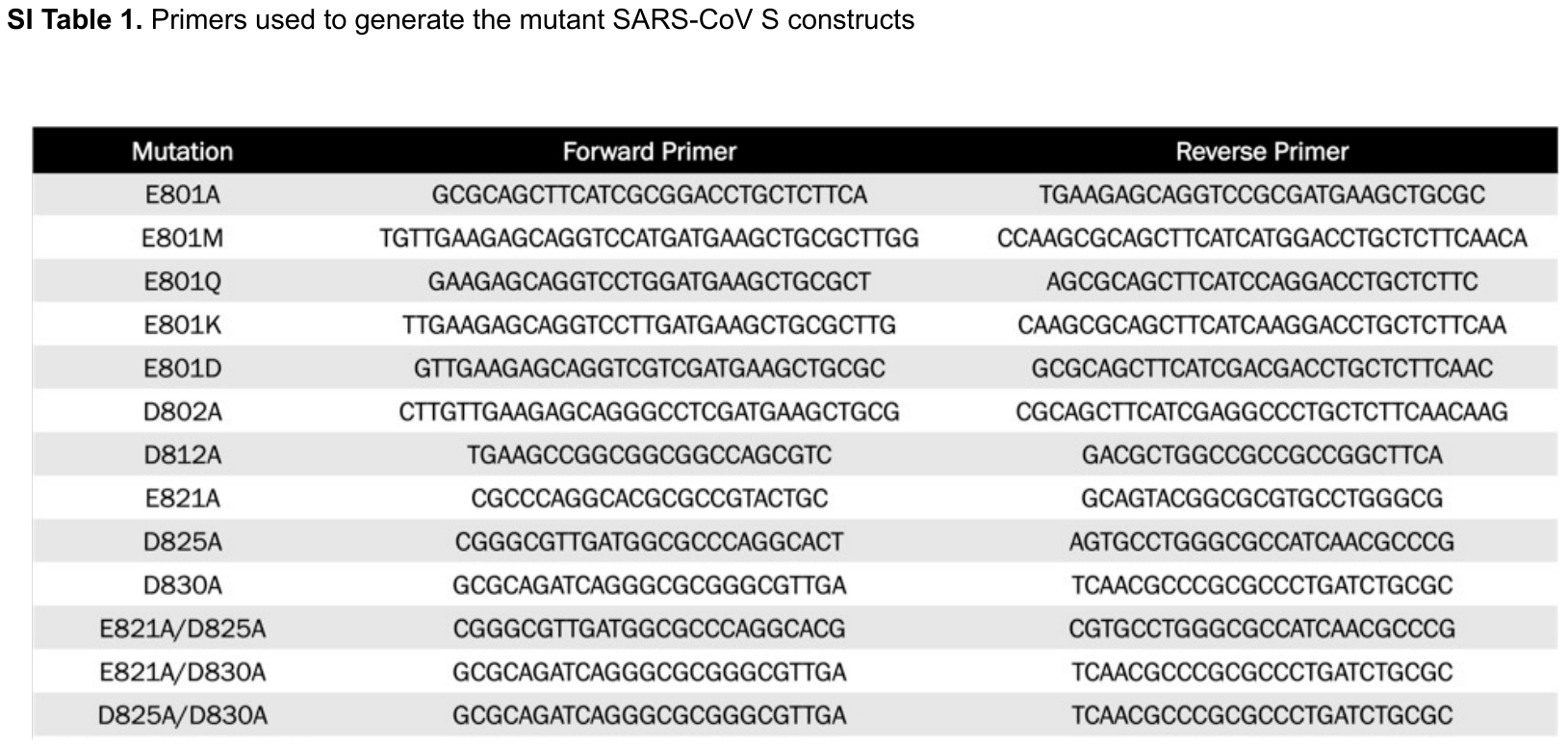
